# Divergent Immune Priming Responses Across Flour Beetle Life Stages and Populations

**DOI:** 10.1101/052472

**Authors:** Imroze Khan, Arun Prakash, Deepa Agashe

## Abstract

Growing evidence shows that low doses of pathogens may prime the immune response in many insects, conferring subsequent protection against infection in the same developmental stage (within life stage priming), across life stages (ontogenic priming), or to offspring (trans-generational priming). Recent work also suggests that immune priming is a costly response. Thus, depending on host and pathogen ecology and evolutionary history, tradeoffs with other fitness components may constrain the evolution of priming. However, the relative impacts of priming at different life stages and across natural populations remain unknown. We quantified immune priming responses of 10 natural populations of the red flour beetle *Tribolium castaneum*, primed and infected with the natural insect pathogen *Bacillus thuringiensis*. We found that priming responses were highly variable both across life stages and populations, ranging from no detectable response to a 13-fold survival benefit. Comparing across stages, we found that ontogenic immune priming at the larval stage conferred maximum protection against infection. Finally, we found that various forms of priming showed sex-specific associations that may represent tradeoffs or shared mechanisms. These results suggest that sex-, life stage-, and pathogen-specific selective pressures can cause substantial divergence in priming responses even within a species. Our work highlights the necessity of further work to understand the mechanistic basis of this variability.

## INTRODUCTION

Immunologists have long assumed that insects lack immune memory and specificity because they do not have the lymphocytes and functional antibodies that are responsible for acquired immunity in vertebrates (Janeway & Medzhitov, 2002). However, growing evidence suggests that a low dose of a pathogen may prime the immune response in insects, reducing the risk and severity of infection by the same pathogen later in life. Evidence for such priming-induced immune protection has been reported in many insects including mealworm beetles (Daukšte *et al.*, 2012), bumble bees (Sadd & Schmid-Hempel, 2006; Tidbury *et al.*, 2011), silkworms (Miyashita *et al.*, 2014), fruit flies (Pham *et al.*, 2007), mosquitoes (Contreras-Garduño *et al.*, 2014) and flour beetles (Roth *et al.*, 2009). Immune priming can also confer sustained protection via (A) ontogenic priming, where the benefit of priming can persist through metamorphosis (Thomas & Rudolf, 2010; Moreno-García *et al.*, 2015) and (B) trans-generational immune priming, where the benefits are manifested in the next generation (Sadd & Schmid-Hempel, 2006; Sadd & Schmid-hempel, 2009; Moreau *et al.*, 2012; Zanchi *et al.*, 2012; Dubuffet *et al.*, 2015). Theoretical models show that within- and trans-generational immune priming can significantly alter pathogen persistence (Tidbury *et al.*, 2012) and reduce infection intensity in populations (Tate & Rudolf, 2012). Thus, it is clear that immunological memory is widespread in insects, and immune priming may have large impacts on the outcome of host-pathogen interactions.

Although we have begun to understand immune priming in many insects, it is not clear how priming evolves. This is partly because the strength, consistency and relevance of immune priming in natural populations remains largely unexplored and is difficult to gauge from laboratory studies. Other aspects of immune function (post-infection survival and encapsulation ability) vary across fruit fly populations (Kraaijeveld, 1995; Corby-Harris & Promislow, 2008), and parasite burden is strongly correlated with the strength of the innate immune response across damselfly populations (Kaunisto & Suhonen, 2013). Similarly, immune priming responses may also vary across natural populations. In laboratory populations, immune priming is affected by the presence of other pathogens (Sadd & Schmid-hempel, 2009) and food availability (Freitak *et al.*, 2009). However, the impact of these factors on immune priming in natural populations is unknown. Wild populations likely face substantial spatial and temporal variation in pathogen diversity, pathogen abundance, and resource availability, generating variability in the strength of selection on immune priming. Priming also imposes fitness costs in some laboratory populations (Contreras-Garduño *et al.*, 2014), potentially generating tradeoffs with other immune responses, or between different types of immune priming. Finally, these fitness costs may also vary as a function of sex and developmental stage. For instance, life-history theory predicts that females should generally evolve higher immune competence than males (Rolff, 2002; Nunn *et al.*, 2009); hence, males may gain more benefits from priming than females (Moreno-García *et al*., 2015). Similarly, variable costs of infection across life stages are also predicted to select for stronger priming responses at specific developmental stages (Tate & Rudolf, 2012). A detailed analysis of such variability can indicate factors that influence the evolution of immune priming. Unfortunately, very few studies have quantified priming in wild insect populations (but see (Reber & Chapuisat, 2012) (ants), (Gonzalez-Tokman *et al.*, 2010) (damselflies), and (Tate & Graham, 2015) (closely related flour beetle species)), and none have measured variation in priming responses across multiple natural populations.

We systematically analyzed immune priming responses of 10 populations of the red flour beetle *Tribolium castaneum* collected from different locations across India (Fig S1). In the laboratory, flour beetles show within life stage (WLS) (Roth *et al.*, 2009), ontogenic (ONT) (Thomas & Rudolf, 2010) and trans-generational (TG) immune priming (Roth *et al.*, 2010), making them an ideal model system to understand the occurrence and abundance of these different types of immune priming responses. We addressed three major questions: (a) Does the immune priming response vary across natural populations and as a function of sex and life stage? (b) Are the different types of priming responses equally beneficial? (c) Are the different types of immune priming responses correlated? Our work is the first report of large within-species variability of priming response across sexes and life stages in natural insect populations. We found that ontogenic immune priming provided greater protection against re-infection, compared to within life stage or trans-generational priming. Finally, our data reveal novel sex-specific links between various forms of immune priming, perhaps representing tradeoffs or even shared mechanistic basis. We hope that our results motivate further investigations to confirm and understand the ecological, evolutionary and mechanistic basis of the observed variability and associations between priming at different stages.

## METHODS

### Beetle collection and experimental individuals

Although immune priming responses should be measured on individuals directly collected from the wild (i.e. grain warehouses), this is difficult to do for the following reasons. First, natural beetle populations do not always have enough individuals of different stages to allow sufficient replication. Second, it is impossible to account for the many factors that may increase within-population variability in immune responses, such as individual age, migration and diet history, and immediate local environment. Controlling for within-population variability in immune priming is essential to quantify variability between populations, which was the major goal of our study. Hence, we established large laboratory populations using wild-collected beetles (maintaining most of the initial genetic variability), and then quantified the immune priming response of individuals of the same age reared under identical conditions. We collected 50-100 *T. castaneum* adults from a grain warehouse in each of 9 cities across India. Of the 10 populations analyzed here, 8 were from different cities and 2 were collected from different warehouses in a single city (Fig S1). We allowed all adults from a site to oviposit for a week on whole-wheat flour at 34°C to start a large laboratory population (>2000 individuals). We maintained these stock populations on a 45-day discrete generation cycle for 9-10 generations before starting experiments.

To generate experimental individuals of equivalent age from all populations, we allowed ~1000 adults from each population to oviposit in 350 g wheat flour for 48 hours. We removed the adults and allowed offspring to develop for ~3 weeks until pupation, collecting pupae daily after this period. We housed 3-4 pupae of each sex separately in 2 ml micro-centrifuge tubes containing 1 g flour for 2 weeks. Since pupae typically eclose in 3-4 days, we obtained ~11-day-old sexually mature virgin adults for immune priming experiments. For experiments with larvae, we allowed adults to oviposit in 350 g flour for 24 hours and collected larvae after 10 days (eggs hatch in 2-3 days; thus, experimental larvae were ~8 days old). In a separate experiment, we found that eggs from all populations developed at a similar rate (Fig S2), confirming that we tested all populations at equivalent developmental stages.

### Immune priming and challenge

For each type of immune priming, we tested all populations together to allow a direct comparison across populations. However, given logistical constraints, we had to test males and females in separate blocks. Note that we only measured maternal TG immune priming in our experiments, and did not measure paternal TG priming. The timeline for each type of immune priming is given in Fig 1 (see supplementary information for detailed methods). For all infections, we used a strain of *Bacillus thuringiensis* (DSM. No. 2046). Originally isolated from a Mediterranean flour moth, this is a natural insect pathogen that imposes significant mortality in flour beetles (Abdel-Razek *et al.*, 1999). On the evening before priming, we inoculated 10 ml nutrient broth (Difco) with cells from a −80°C stock of *B. thuringiensis*. We incubated the growing culture overnight in a shaker at 30°C until it reached an optical density of 0.95 (measured at 600 nm in a Metertech UV/Vis Spectrophotometer, SP8001). We centrifuged the culture at 5000 rpm for 10 minutes, removed the supernatant, and resuspended the pellet in 100μl insect Ringer solution (7.5g NaCl, 0.35g KCl, 0.21g CaCl_2_ per liter) to make bacterial slurry. We killed the bacteria in a heat block at 90°C for 20 minutes as described earlier (Roth *et al.*, 2009; Khan *et al.*, 2015). We used heat-killed bacteria to prime individuals, since this would elicit an immune response without any direct cost of infection.

**Figure 1.**
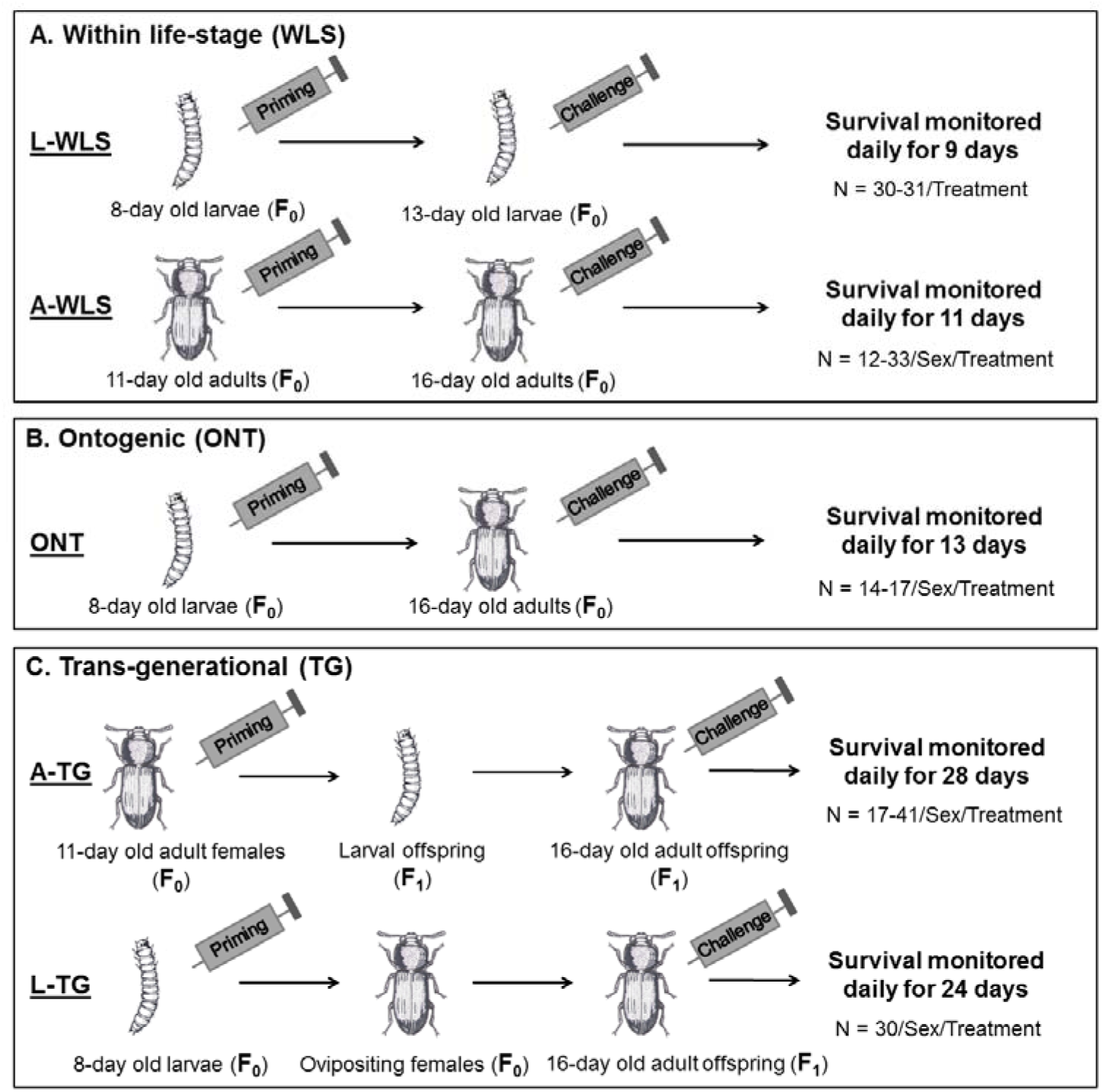
Experimental design to measure the strength of immune priming responses at different stages: (A) Within life stage priming (individuals primed and challenged as larvae (L-WLS) or adults (A-WLS)) (B) Ontogenic priming (individuals primed as larvae and challenged as adults) (C) Trans-generational maternal priming (females primed as larvae (L-TG) or adults (A-TG) were paired with uninfected virgin males and their offspring were challenged). Sample sizes are indicated for each treatment (priming and control) and sex.

To prime individuals, we pricked them with a 0.1 mm minuetin pin (Fine Science Tools, Fosters City, CA) dipped either in heat-killed bacteria (primed) or in sterile insect Ringer solution (control). To minimize damage to internal organs we pricked individuals laterally between the head and thorax (adults) or between the last two segments (larvae). After priming (or mock priming), we isolated individuals in wells of 96-well microplates containing flour. When appropriate, we sexed pupae and distributed them individually in wells of 96-well microplates. For subsequent immune challenge, we pricked individuals as described above, but used live bacterial slurry (without heat-killing). After this, we again isolated individuals in fresh microplates and monitored their survival (See Fig 1 for timeline).

### Data analysis

We analyzed post-infection survival data for each population, sex and immune priming type separately using Cox Proportional Hazard survival analysis with priming treatment as a fixed factor (see Figs S3-S11 for survival curves). We noted individuals that were still alive at the end of the experiment as censored values. We calculated the strength of a given type of immune priming response within each population (and sex) as the estimated hazard ratio of unprimed vs. primed groups (hazard ratio = rate of deaths occurring in unprimed group/ rate of deaths occurring in primed group). A hazard ratio significantly greater than one indicates a greater risk of death after infection in the unprimed (control) compared to primed individuals.

To estimate the overall impact of sex on the immune priming response, we analyzed hazard ratios using a two-way ANOVA with sex and type of immune priming as fixed factors. We excluded data from larval within life stage priming (L-WLS) because sex cannot be distinguished in larvae. To test whether the strength of the priming response varies as a function of life stage at priming (larvae vs. adults), we analyzed hazard ratios with a one-way ANOVA. Finally, to compare the strength of priming across different stages (Fig 1), we analyzed data with a one-way ANOVA and used Tukey’s honest significant difference (HSD) to estimate pairwise differences after correcting for multiple comparisons.

We also wanted to test whether the strength of immune priming responses was correlated across types of priming. However, several populations did not show a significant immune priming response; hence, we could not use a linear regression approach. Therefore, we generated a contingency table, categorizing each population according to the presence (proportional hazard test: p < 0.05) or absence (proportional hazard test: p > 0.05) of each type of priming response (also see Figs S3-S11). We then used a Fisher’s exact test to determine whether the presence of the two types of immune priming was qualitatively associated across populations.

## RESULTS

### The immune priming response varies across populations

We estimated the strength of immune priming as the proportional hazard ratio of individuals mock-primed with sterile Ringer solution vs. primed with a pathogen (heat-killed *B. thuringiensis*), followed by a subsequent infection with live *B. thuringiensis*. Surprisingly, we found that only about half the populations showed significant priming at a given stage, although all populations were capable of mounting multiple forms of immune priming (Fig 2). The immune priming response varied substantially in larvae as well as adult males and females across natural populations (Fig 2; Figs S3-S11). We found that only a few populations showed significant within life stage immune priming as larvae (L-WLS, 4/10 populations) or as adults (only females; A-WLS, 4/10 populations) (Fig 2A). In contrast, at least one sex of many populations showed significant ontogenic (ONT, 9/10 populations; Fig 2B) and trans-generational benefits of adult priming (A-TG, 6/10 populations; Fig 2C). Our data also demonstrate long ranging impact of trans-generational immune priming in several populations (L-TG, 6/10 populations; Fig 2D), whereby priming larvae improved post-infection survival of their adult offspring. Finally, we found that populations B1 and B2 showed very different priming responses (Fig 2), although they were collected from different warehouses in the same city. Hence, geographical proximity does not seem to be a good predictor of similarity in immune responses.

**Figure 2.**
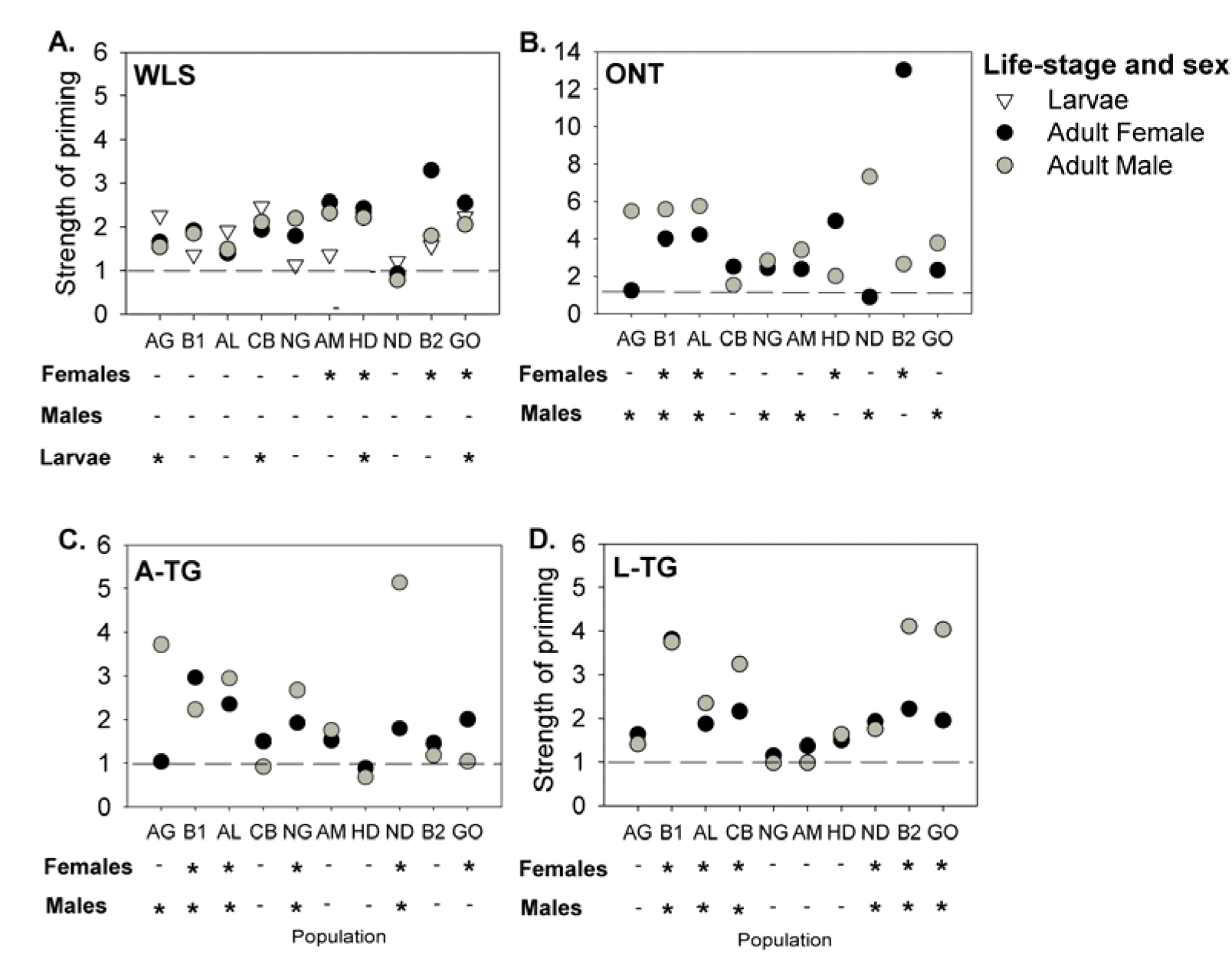
Variation in priming response across sexes, life stages and populations. (A) Within life stage immune priming (WLS) benefit in larvae and adults (B) Ontogenic (ONT) immune priming benefit (C) Trans-generational (TG) immune priming benefits from adult females (D) Trans-generation (TG) immune priming benefits from larvae. Strength of immune priming response was calculated as the hazard ratio of the proportion of deaths occurring in the unprimed group compared to the primed group under proportional hazard model. Horizontal dashed lines in each panel indicate a hazard ratio of 1. ‘*’ and ‘-’ denote significant (p ≤ 0.05) and nonsignificant (p > 0.05) impact of immune priming in each stage, sex, and population. Sample sizes for each group are given in Fig. 1.

### Effect of sex on immune priming

As explained in the methods, we tested the priming response of each sex separately. Hence, we could not directly test for an impact of sex in each population. Combining hazard ratios across populations, we did not find a consistent impact of sex on the strength of the immune priming response for any type of priming (Table 1A-C). However, in many populations, only one sex showed a significant priming response. For instance, the adult WLS response appears to be female-limited, with males showing no priming in any population (Fig 2A). Similarly, in most populations that showed ontogenic priming, priming was beneficial for only one sex (7/9 populations; Fig 2B). However, unlike WLS, we did not find a systematic benefit of ONT priming: the sex that benefited from ONT priming varied across populations. We also failed to find clear sex-specific benefits of TG priming for offspring. We observed adult maternal immune priming (A-TG) in offspring of both sexes (4 populations) or only one sex (2 populations) (Fig 2C). Intriguingly, all six populations with significant larval trans-generational (L-TG) priming showed a response in offspring of both sexes (Fig 2D). Thus, both males and females tend to show parallel benefits of L-TG priming across populations. Overall, our results show that the impact of sex on immune priming varies both across populations and type of immune priming.

**Table 1.** Summary of (A) two way ANOVA for immune priming response with type of immune priming and sex as fixed factors (B) one way ANOVA for WLS and ONT priming response with sex as a fixed factor (C) two way ANOVA for TG priming response with sex and type of parental priming (e.g. larval or adult priming) as fixed factors (D) one way ANOVA for immune priming response with life stage-specific (larvae or adults) priming as a fixed factor (E) one way ANOVA for immune priming response with type of immune priming response as a fixed factor. IP = Immune priming, S = Sex, PP = Type of parental priming, LS = Life stage.

| Experiment | Effect | df | SS | F-ratio | P |
| --- | --- | --- | --- | --- | --- |
| **A. Impact of type of IP and S** | IP | 2 | 4.632 | 9.24 | **<0.001** |
| (excluding L-WLS) | S | 1 | 0.126 | 0.503 | 0.48 |
|  | IP × S | 2 | 0.268 | 0.536 | 0.587 |
| **B. Impact of S on A-WLS response** | S | 1 | 0.044 | 0.37 | 0.54 |
| Impact of S on ONT response | S | 1 | 0.249 | 0.61 | 0.44 |
| **C. Impact of S and PP on TG** | S | 1 | 0.148 | 0.601 | 0.443 |
| response | PP | 1 | 0.167 | 0.678 | 0.415 |
|  | S×PP | 1 | 0.000 | 0.003 | 0.951 |
| **D. Impact of priming at larvae vs. adults** | LS | 1 | 1.58 | 5.99 | **0.016** |
| **E. Impact of type of IP (including L-WLS)** | IP | 4 | 5.23 | 5.67 | **<0.001** |

### Larval ontogenic priming maximizes protection against subsequent infection

Next, we tested the impact of priming life stage on the strength of the priming response. We found that priming at the larval stage was more beneficial and produced a greater response than priming adults (Table 1D). However, this result was driven primarily by ontogenic larval priming, which maximized post-infection survival in adults across priming types relative to the respective unprimed controls (Fig 3, Table 1E). Larval ONT priming resulted in a ~3 fold survival benefit, compared to the 2-fold benefit observed for other forms of priming, including larval WLS priming (Fig 3). We also found that across populations, the strength of ONT priming in females was more variable compared to WLS, L-TG or A-TG priming (Bartlett’s test for homogeneity of variance, p < 0.02 for each pairwise comparison; compare boxplots in Fig 3). For males, ONT priming was significantly more variable than WLS priming, but not other forms of priming. Together, our results suggest that among different types of immune priming, ONT priming responses are strongest and most variable.

**Figure 3.**
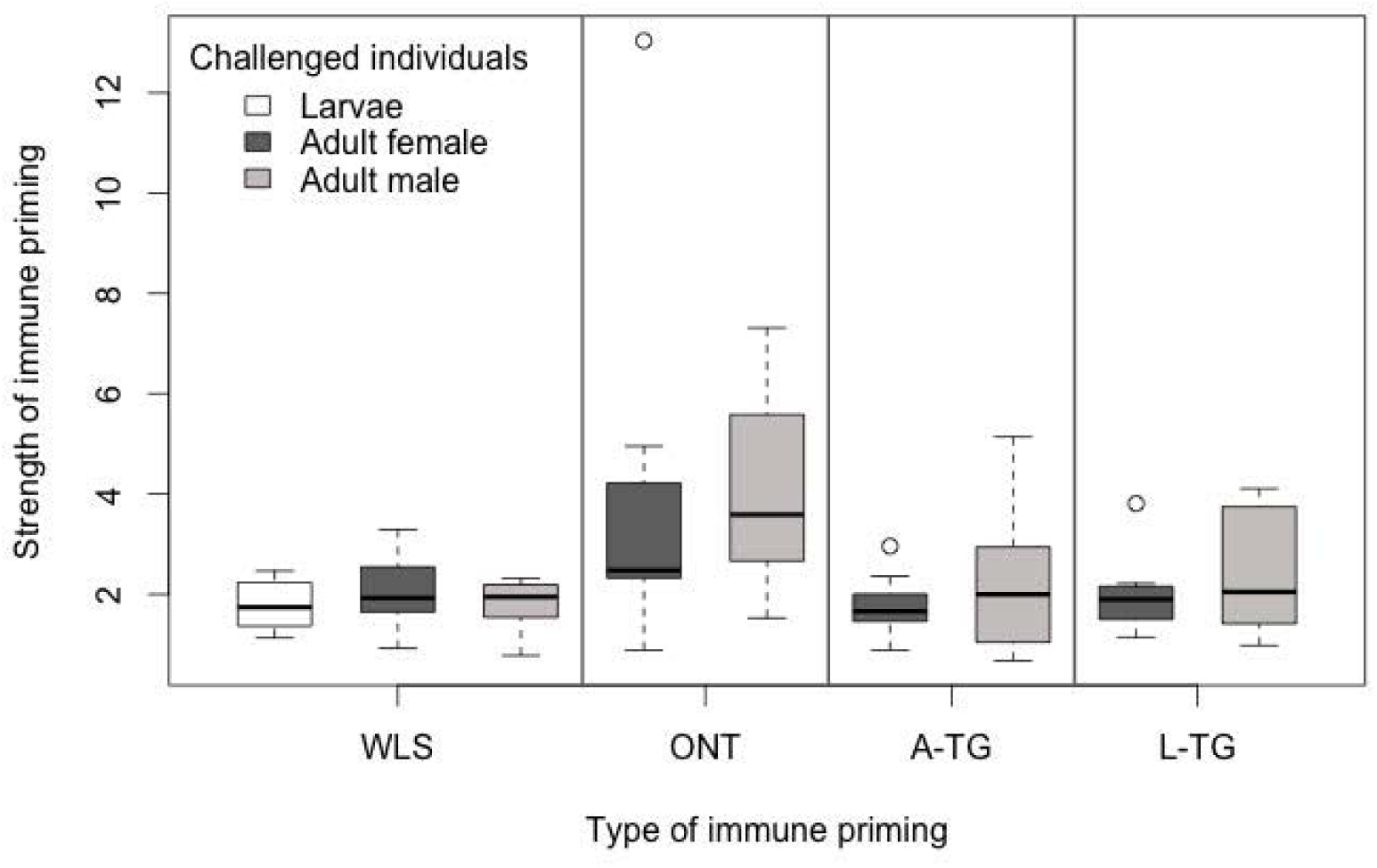
Strength of each type of immune priming response across different life stages and sexes. Strength of priming was calculated as described in Fig. 2. Sample sizes for each assay are shown in Fig. 1. WLS = within life stage immune priming, ONT = ontogenic priming; A-TG = trans-generation benefits of adult (maternal) priming; L-TG = trans-generation benefits of larval priming.

### Associations between within- and trans-generation immune priming

We tested whether different types of immune priming responses were associated within populations. We found that most populations either showed significant female WLS priming or significant TG priming in male offspring, but not both (Fig 4A; Fisher’s exact test, p = 0.046). In contrast, there was no association between female WLS and TG priming in female offspring (Fig S12A; Fisher’s exact test, p = 0.643). We also found a non-significant trend for an association between ONT priming in males and TG priming in male offspring (Fig 4B; Fisher’s exact test, p = 0.446), but not for female offspring (Fig S12B; Fisher’s exact test, p = 0.663). For male offspring, one of the two populations that showed only ONT priming had nearly significant TG priming (population AM, Fig 4B; p = 0.066). If this population were counted as showing both types of priming, the association between ONT and male TG priming would be significant (Fisher’s exact test, p = 0.046). Although the association is not strong, these results suggest that in populations where male adults benefit from larval ONT priming, they may also benefit from maternal TG immune priming. Overall, our results indicate that trans-generational immune priming responses are associated with within-generation responses, but the association is limited to male offspring.

**Figure 4.**
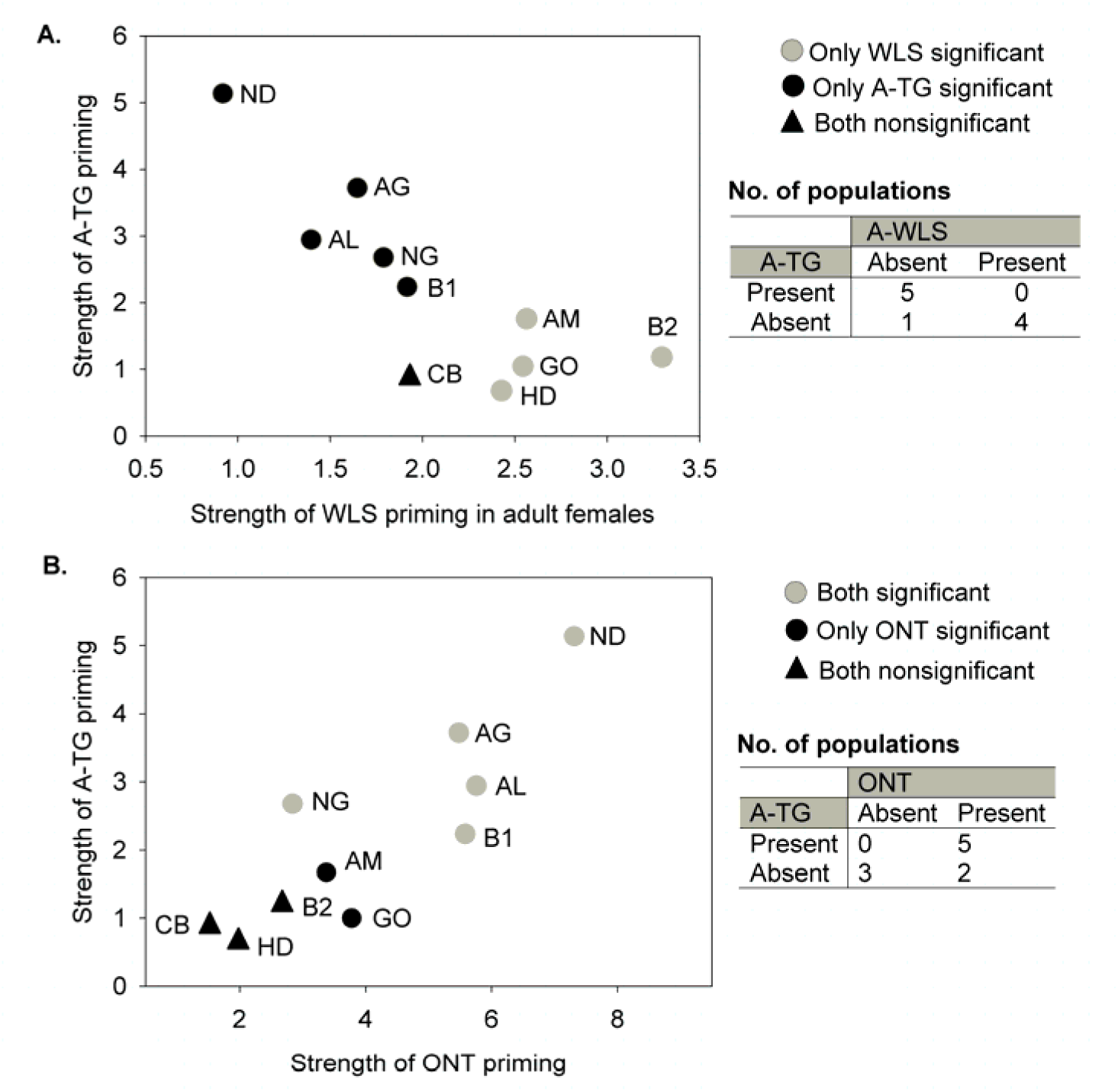
Associations between within- and trans-generation immune priming. Strength of A-TG response in male offspring as a function of (A) strength of WLS immune priming in female adults (B) ONT priming in males. Strength of priming was estimated as described in Fig 2. Each population (labelled) was categorized based on the presence or absence of each type of priming response (using significant hazard ratios as explained in Fig 2), and contingency tables (shown beside each panel) were used to test the association between two types of immune priming across populations. WLS = Within life stage immune priming, A-TG = Trans-generational benefits of adult (maternal) priming, ONT = Ontogenic immune priming.

## DISCUSSION

Our work provides the first evidence of substantial variation in both within- and trans-generational immune priming responses among natural populations of an insect. Approximately half the populations did not show a significant response to any given type of priming; on the other hand, all populations showed at least two forms of priming. Relative to unprimed controls, primed individuals showed up to 13-fold higher survival in some cases, whereas others showed no benefits of priming. Note that we reared wild-collected beetles under standard laboratory conditions for 9-10 generations before starting our experiments; hence, we probably underestimated the variation in priming responses across populations. What is the cause of this variability? Potential hypotheses include gain and loss of priming responses via genetic drift; local adaptation to specific pathogen diversity and abundance (Sutton *et al.*, 2011); variable life-history related costs associated with immune investment (Roy & Kirchner, 2000; Miller *et al.*, 2006); and variable susceptibility to pathogens (Best *et al.*, 2013). Currently, we cannot directly test these hypotheses since we do not have information on the local pathogen pressure experienced by our beetle populations, the fitness costs of immune priming, or their relative susceptibility to *B. thuringiensis*. Nonetheless, our work demonstrates the importance of quantifying variability of immune priming responses in natural populations, and sets up a framework to understand the evolution of immune priming responses.

One of our most interesting findings is that ontogenic priming confers a greater survival benefit than within life stage or trans-generational immune priming response. A recent theoretical model predicts that if adults incur higher costs of infection than larvae, selection should favor strong ontogenic priming that reduces the proportion of susceptible adults (Tate & Rudolf, 2012). On the other hand, trans-generational priming should be favored when larvae are more susceptible to infection than adults. Thus, if *B. thuringiensis* imposes stage-specific costs of infection in *T. castaneum*, it may have selected for stronger ontogenic priming in our populations. In a separate experiment, we found that larvae and adults from a laboratory-adapted, outcrossed flour beetle population were equally susceptible to *B. thuringiensis* infection (Fig S13A). These data suggest that beetle life stages are not differentially susceptible to infection, although it is possible that our natural populations do show stage-specific susceptibility. Another interesting result from our analysis is that the strength of larval TG priming is similar to the strength of adult TG priming, but much weaker than larval ontogenic priming. Thus, the high survival benefit of ONT priming (through metamorphosis) is not transmitted to the next generation. Thus, we speculate that during oviposition, priming is “reset”, perhaps because the mechanisms responsible for ontogenic and trans-generational priming are different. Further empirical studies are thus critical to elucidate the complex interplay between immune priming types and their relative impact on the outcome of infection within a population.

Our data also revealed novel associations between within- and trans-generational immune priming responses. In populations where adult females showed significant within life stage immune priming, male offspring did not show trans-generation priming. We speculate that this negative relationship may reflect a trade-off between maternal and offspring immunity (Moreau *et al.*, 2012): transferring immunity to offspring may be costly for females who also bear the cost of their own immune priming response. However, this needs to be explicitly tested by quantifying the difference in the priming response of offspring of individual females that were primed and challenged as adults, vs. females that were not primed and challenged. Our results also suggest a weak association between male ONT and male TG priming. Interestingly, both relationships between trans-generational and within-generation priming were limited to male offspring. Such male-specific associations may arise due to sex-specific variation in infection susceptibility, investment in other immune components, or tradeoffs with other fitness components. We cannot test these predictions since the relative impact of *B. thuringiensis* infection in both sexes is unknown in natural beetle populations. However, separate experiments with an outbred *T. castaneum* population showed that infected males die about twice as fast as females (Fig S13B). It is possible that the natural populations analysed here also show similar sex-specific variation in susceptibility to infection, and further work is necessary to distinguish between these hypotheses.

We suggest that our results are applicable in many insect-pathogen systems. *B. thuringiensis* infects multiple insect hosts (Bravo *et al.*, 2011), and is commonly found in diverse habitats such as soil, insect cadavers, water and grain dust (Argôlo-filho & Loguercio, 2014; Lambert & Peferoen, 2014). Hence, *B. thuringiensis* may impose strong selection on many insects occupying diverse ecological niches, influencing the evolution of their immune responses in the wild. Although we did not test whether the immune priming response is specific to the *B. thuringiensis* strain that we used, an earlier study showed that *T. castaneum* individuals could differentiate between strains of the same pathogen (Roth *et al.*, 2009). Thus, the immune priming response that we observed is most likely a specific response against *B. thuringiensis* and does not represent general protection via an overall upregulation of immune components. Finally, we assayed immune priming response using septic injury, whereas many pathogens infect their insect hosts via the oral route. However, recent studies confirm that both septic injury (Roth *et al.*, 2009) and oral infection (Milutinović *et al.*, 2014) with *B. thuringiensis* produce comparable immune priming responses in *Tribolium* beetles, suggesting that our infection protocol is unlikely to bias our results.

We would like to end by highlighting several open questions that have emerged from our work. (A) Do sex- and stage-specific differences in immune function and pathogen susceptibility explain the observed variation in immune priming response? (B) Do variable fitness costs of immune priming explain the observed variation in immune priming response across populations? (C) Finally, do mechanisms underlying various forms of immune priming differ from each other? We suggest that future work on insect immune priming should focus on variation in both the mechanistic as well as ecological and evolutionary aspects of natural variation in immune priming. In particular, experimental manipulation of specific immune priming types across sexes and life stages promises to shed light on the complex problem of immune priming responses and their variable outcomes in natural populations.

## COMPETING INTERESTS

We have no competing interests.

## AUTHOR CONTRIBUTIONS

IK and DA conceived of and designed experiments; IK and AP carried out experiments; IK and DA analyzed data; DA and IK wrote the manuscript with input from AP. All authors gave final approval for publication.

## ACKNOWLEDGEMENTS

We thank members of the Agashe lab for critical comments on the manuscript, and NG Prasad for the *B. thuringiensis* strains.

## FUNDING

We acknowledge funding and support from a SERB-DST Young Investigator Grant to IK, a DST INSPIRE Faculty fellowship to DA, and the National Center for Biological Sciences (NCBS), India.

